# Effect of Acute Stress on Working Memory in Pilots: Investigating the Modulatory Role of Memory Load

**DOI:** 10.1101/2023.06.23.546263

**Authors:** Yaowei Liang, Xing Peng, Yueying Liu, Qi Zhu, Zhi Xu, Jiazhong Yang

## Abstract

The effect of acute stress on working memory (WM) has not been extensively investigated in demanding tasks, although many practitioners, such as pilots, frequently face varying working memory demands. The current study investigated how acute stress modulates pilots’ WM performance. Forty-two pilots were randomly assigned to the stress group or the control group. The stress group experienced acute stress induced by the modified Trier Social Stress Test (TSST), whereas the control group did not receive any stress induction. Both groups performed N-back tasks under varying degrees of memory load (0-back, 1-back, and 2-back). State-Trait Anxiety Inventory scores, heart rates, and salivary cortisol concentrations were measured in the experiment to determine the extent of induced stress. The results revealed that (1) The modified TSST effectively induced acute stress in the stress group. (2) As rising of memory load, the reaction time expectedly increased while accuracy decreased. (3) An interaction between the stress and control group was found for accuracy, indicating that acute stress might contribute to higher accuracy in moderate WM load (1-back). These suggestions support the emotion regulation hypothesis, in which acute stress induces heightened emotional states that impact WM performance. Furthermore, the current study explored the interplay between acute stress and memory load in a specific population, offering insights into potential pilots’ stress management strategies and training programs to improve pilots’ WM abilities. These applications would be beneficial for strengthening pilots’ coping mechanisms in emergencies and guaranteeing flight safety.

## Introduction

Working memory is a cognitive system crucial for the temporary storage and manipulation of information [1], which plays an essential role in complex flight operations. In the highly demanding field of aviation, pilots frequently face unavoidable and acute stress because of weather conditions, technical malfunctions, and air traffic congestion. Acute stress caused by these factors has the potential to impair cognitive functions such as working memory (WM) [2, 3] through brief periods of psychophysiological and behavioral changes. Therefore, understanding the relationship between acute stress and WM is crucial for flight safety and performance.

The Trier Social Stress Test (TSST) is a well-established paradigm used to induce stress [4]. It consists of a videotaped free speech and a subsequent mental arithmetic task for 15 minutes in front of a nonresponsive audience. It is an effective method to raise physiological and psychological responses to acute stress. In the physiology field, stress is mainly related to the fast response pathway of the Sympathetic Nervous System (SNS), manifesting as an increased heart rate. Slow-response pathways were involved in activating the Hypothalamic Pituitary Adrenal (HPA) axis [5, 6] in terms of increased cortisol secretion.

The N-back task is the classical paradigm for measuring WM [7]. Participants are asked to remember the stimulus presented N items before the current one, with the ‘N’ value manipulating the level of memory load. In aviation activities, WM plays a fundamental role in processing and storing information relevant to ongoing tasks, making it essential for pilots to effectively tackle complex flight operations and respond to emergency situations. This ability to execute these operations methodically is also called Standard Operating Procedures (SOPs), which were found to be affected by WM. For example, the Little Rock air disaster in June 1999 in the U.S. was due to the impaired WM of pilots under acute stress conditions, which caused the pilots to forget SOPs and consequently operate the spoilers incorrectly.

To date, research regarding the impact of acute stress on WM has yielded mixed findings [8-10]. While the consensus is that acute stress can impair WM [11], some studies propose that certain levels of stress may enhance WM performance [12, 13]. For instance, under some stress conditions, individuals have demonstrated improved recall [14]. Further confounding this issue is the potential counteraction of memory load, which has been suggested to account for these discrepancies [15]. Therefore, the specific impact of acute stress on pilots’ WM and the role of memory load in this relationship remain underexplored.

To address these gaps, this research aims to examine the specific effects of acute stress on pilots’ WM and to scrutinize the potential moderating role of memory load. We utilized a modified TSST to induce acute stress in pilots and an N-back task to assess their working memory performance under varying memory load conditions. Moreover, the current study included a control group of pilots who did not undergo acute stress, allowing for a more understanding of the interplay of stress and memory load.

We hypothesized that as the memory load increased, both reaction time and accuracy would be negatively impacted, irrespective of acute stress induction. At high levels of memory load, we anticipate that the control group would perform with greater accuracy and shorter reaction time than the stress group.

In summary, this study aims to delve deeper into the complex relationship between acute stress and working memory in pilots, focusing particularly on the potential moderating role of memory load. Given the critical importance of working memory in aviation tasks and the potential impairment posed by acute stress, the findings from this research could hold practical significance in improving pilot training, stress management, and overall performance during flight tasks.

## Materials and methods

### Participants

According to G*power calculations[16], a total sample size of at least 28 individuals was required to provide sufficient power to detect a medium-sized effect (f = 0.25) with 80% power and a significance level of α = 0.05. We recruited forty-two healthy male pilots (aged 21–25 years) with normal or corrected-to-normal vision to participate in this study. Since the target population of the current study is pilots, a special occupational group that often faces high-intensity work stress and challenges, we selected male participants, as males are currently a majority in this profession. All participants received commercial aviation licenses from the Civil Aviation Administration of China (CAAC) and logged an average of more than 230 hours of flight in simulators and real aircraft. The participants were randomly assigned to stress and control groups before the experiment, and both groups were matched on flight hours, age, and education, with 21 participants in each group. They were paid a fee after the experiment. The Ethics Committee of the Civil Aviation Flight University of China approved this study. Each participant gave written informed consent. The authors did not have access to information that could identify individual participants after data collection. The participants were involved in the experiment in May 2022.

## Experimental design

### Experimental procedure

The experiment consisted of three parts: the rest, the Trier Social Stress Test (TSST), and the N-back task session (Fig 1). In the initial phase of the experiment, participants arrived at the laboratory and were guided to a separate room, where they sat undisturbed for 10 minutes. They then filled out basic demographic forms and completed both trait and state anxiety questionnaires. Following this, participants in the stress group underwent a modified version of the Trier Social Stress Test (TSST), whereas the control group underwent the same procedure without stress induction. The final component of the session involved the N-back task, designed to assess working memory.

**Fig 1.**
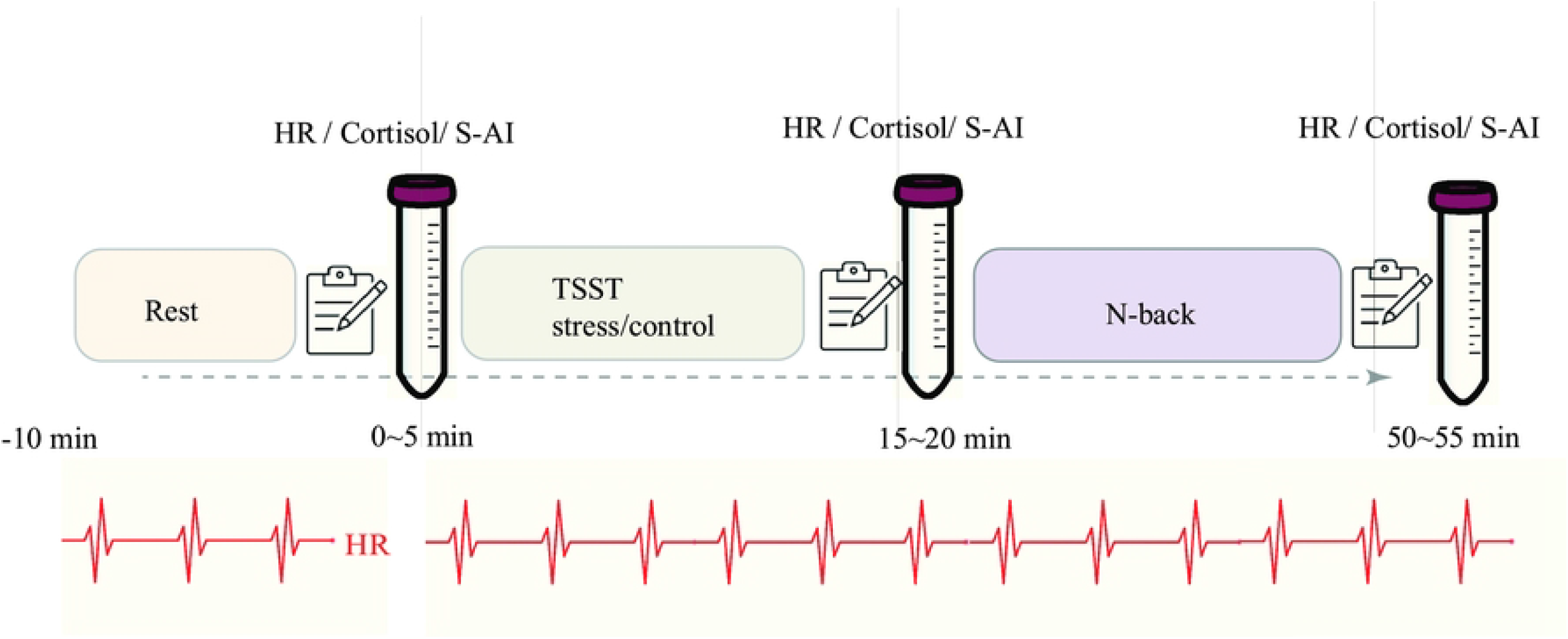
Experimental procedure.

In this study, State_Anxiety Inventory (S-AI) scores, heart rates, and salivary cortisol concentrations were selected as indicators of acute stress, and accuracy and reaction time were used as indicators of WM. Participants were instructed to avoid strenuous exercise and alcohol consumption for 24 hours before the experiment and refrain from eating and smoking for 2 hours before the experiment. S-AI scores, heart rates, and saliva samples were collected three times throughout the experiment: 5 min after the participant’s meditation (baseline), after the TSST, and after the N-back task. The study followed a mixed experimental design with two groups (stress and control) and three levels of memory load (0-back, 1-back, and 2-back), with ‘group’ as a between-subjects variable and ‘load level’ as a within-subjects variable.

### Induction of stress

The TSST requires participants to complete several tasks, including a short preparation time, a videotaped free speech, and a mental arithmetic task [4]. In this study, the modified TSST paradigm was appropriately adapted to the pilots’ characteristics, including preparation time (2 min), an interview (5 min), and a mental arithmetic task (5 min). During the preparation phase, participants could peruse the interview questions. The questions were selected from the oral proficiency interview of the international civil aviation organization (ICAO) and the oral assessment questions of the practical examination, which experts used to form six questions. In the interview phase, participants were randomly selected to answer the questions in front of the experts. In the mental arithmetic phase, participants were asked to report the results quickly and accurately (e.g., starting from 2031, perform the mental arithmetic task in decreasing order of 18). Before the TSST, the stress group was informed that they would be videotaped and scored. Two experts (a psychologist and a flight instructor) were invited to record and evaluate the participant’s performance and posture. The experts wear uniforms and work permits, maintain a neutral expression throughout the process, and make specific reactions when necessary.

Comparable to the stress group, the pilots of the control group participated in a similarly physically and mentally demanding task. In the preparation phase, participants perused material about civil aviation. In the interview phase, participants read the material at a normal speed. In the mental arithmetic task, participants only had to perform simple addition (e.g., mental arithmetic in successive increments of 5 starting from 3). Throughout the entire process, the experts did not wear formal attire, the tasks were not videotaped, and the pilots were not evaluated. It lacked the stress-inducing components of the TSST.

### Working memory test: The numerical N-back task

After the TSST (stress induction vs. control situation), the WM performance of the participants was tested with an N-back task (Fig 2). Participants were conducted in a quiet and dimly lit room, approximately 60 cm from the display. The stimuli were presented on a 17-inch display with a resolution of 1024*768 pixels and a refresh rate of 60 Hz using the software E-prime 2.0. The experimental stimuli were all presented on a black background, and the stimulus was a white digit (e.g., 1∼9). The practice session consisted of 3 blocks of 60 stimulus trials with feedback. It was ensured that participants understood the instructions before the formal experiment. The formal experiment consisted of 3 blocks corresponding to low (0-back), moderate (1-back), and high memory load (2-back), with each block having a total of 180 trials. Of these, 1/3 of the trials were digits that required the participants to perform an “A” key response. The stimulus was presented for 500 ms, followed by a “+” fixation for 1500 ms, for a total response time range of 2000 ms. Participants pressed the “A” and “L” keys with the index finger of their left and right hands, respectively.

**Fig 2.**
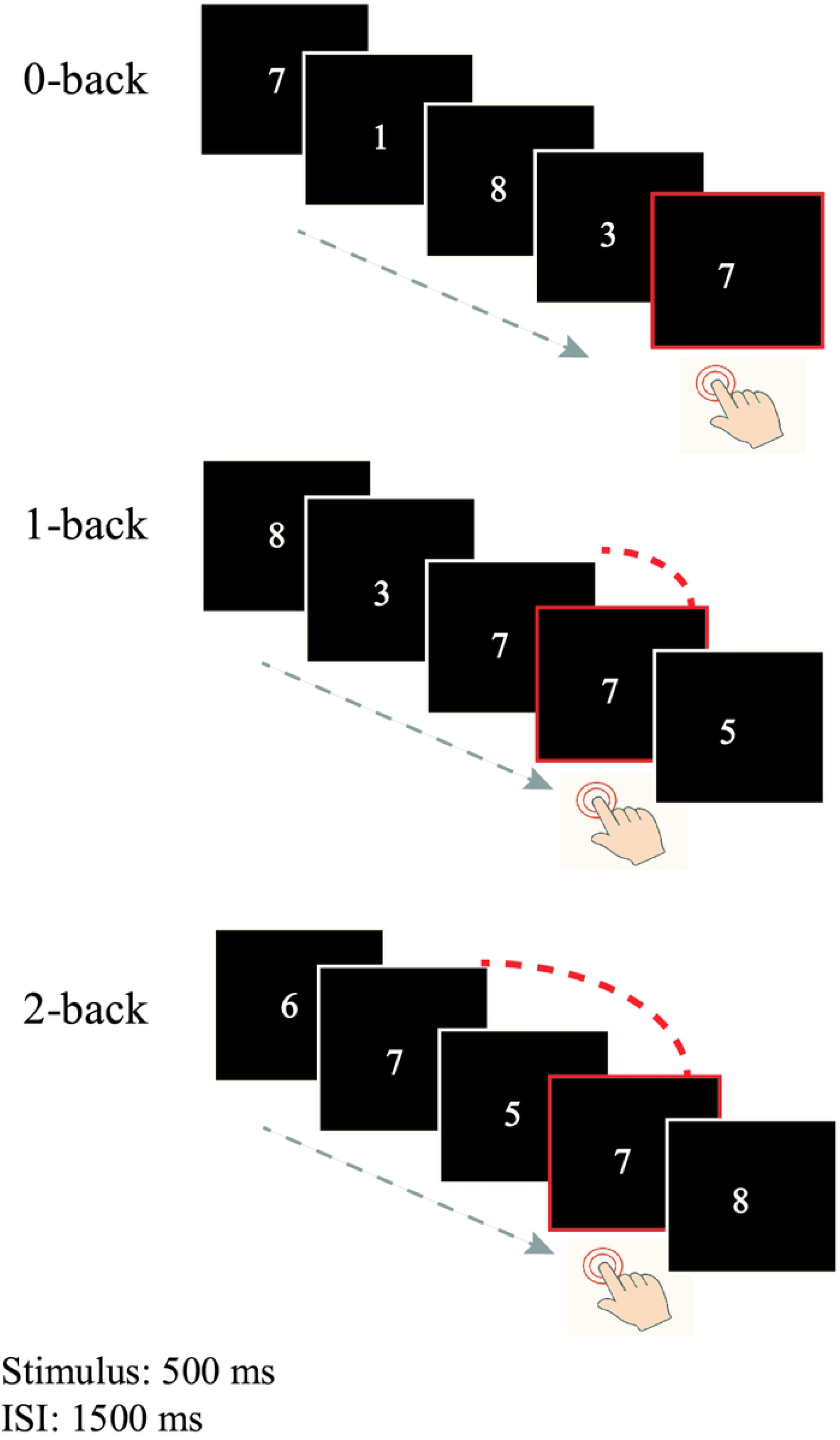
Example of the N-back task with the stimulus material used in the experiments.

In the 0-back condition, participants were asked to determine whether the current stimulus was the same as the first stimulus at the beginning of the experiment, and if it was the same, press key A; if not, press key L. In the 1-back condition, participants were asked to determine whether the current stimulus had appeared one position back in the sequence, and if it was the same, press key A; if not, press key L. In the 2-back condition, participants were asked to determine whether the current stimulus was the same as the second stimulus backward in time, and if it was the same, press key A; if not, press key L.

## Data analysis

Cortisol concentrations and SA-I scores were used as indicators of stress levels. A 2 (group: stress and control groups) × 3 (measurement time points: baseline, post-TSST, and post-N-back task) repeated-measures ANOVA was performed on the data. To calculate the mean accuracy and reaction time for all participants, a 2 (group: stress and control groups) × 3 (memory load: 0-back, 1-back, and 2-back) repeated-measures ANOVA was performed on the data. Greenhouse–Geisser corrected *p* values were used when appropriate.

## Results

### S-AI scores

The change in S-AI scores relative to baseline is shown in Fig 3 (a). There was a significant main effect of group (*F*(1, 40) = 12.35, *p* < 0.001, *η*_*p*_^2^ = 0.24), and the S-AI scores were significantly higher in the stress group than in the control group. The main effect of time was also significant (*F*(2, 80) = 20.76, *p* < 0.001, *η*_*p*_^2^ = 0.34). There was a significant interaction between group and time (*F*(2, 80) = 16.02, *p <* 0.001, *η*_*p*_ ^2^ = 0.29).

**Fig 3.**
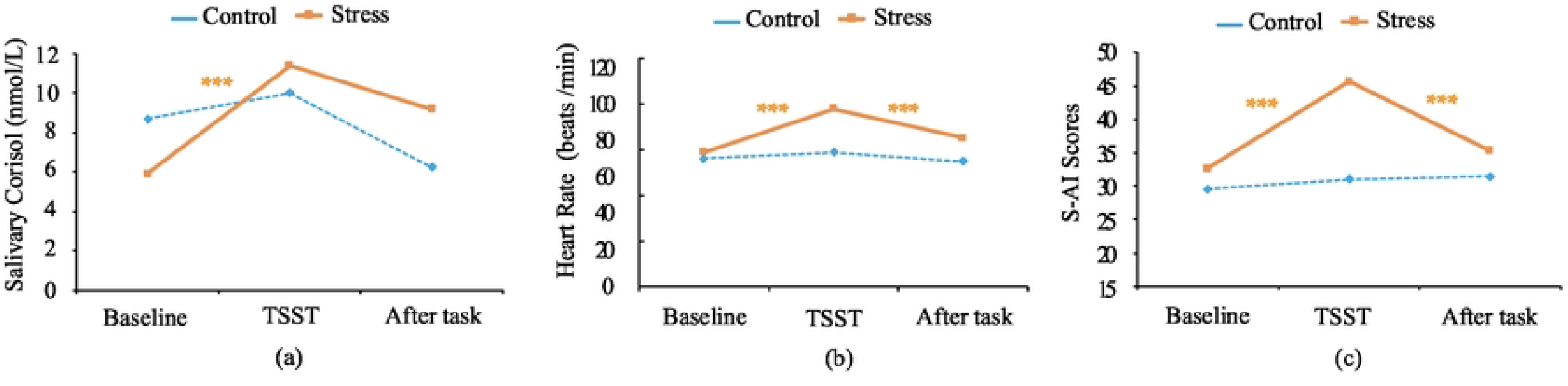
S-AI scores, heart rate (beats/min), and salivary cortisol concentrations (nmol/L) for stress and control pilots at three-time points.

**Fig 4.**
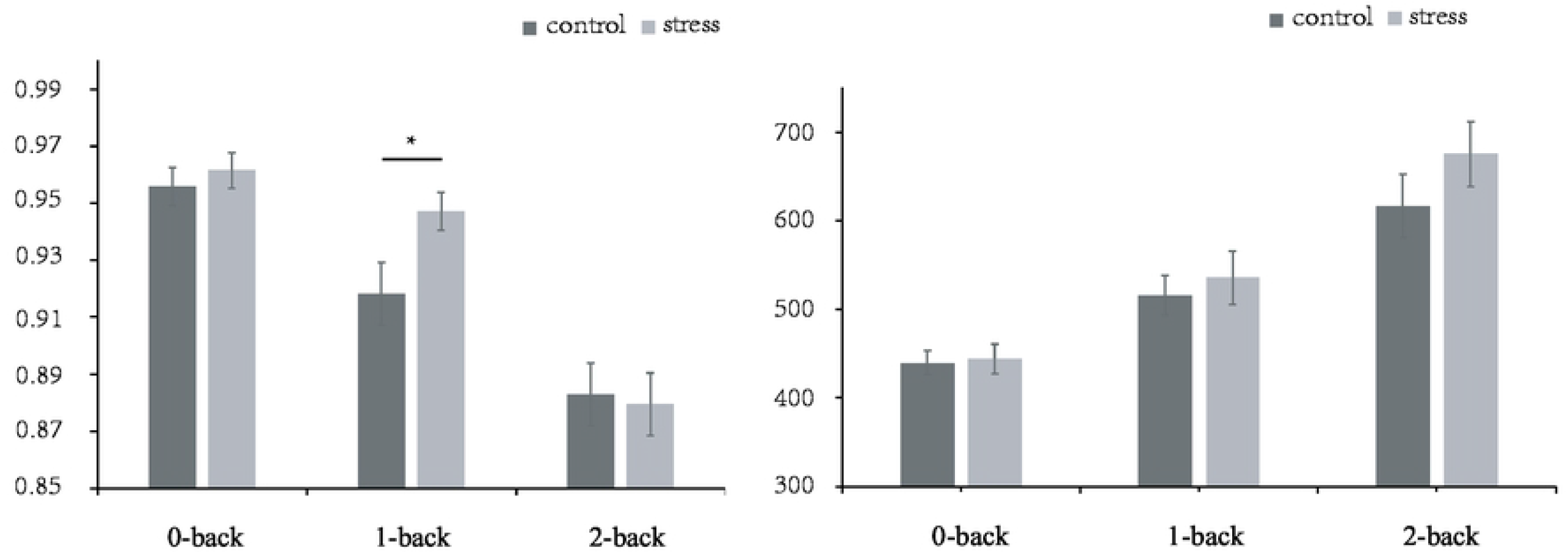
Mean reaction time and accuracy in the 0-back, 1-back, and 2-back conditions.

Simple effect analyses showed that, for the control group, there were no significant differences across the three stages in time: baseline (29.57 ± 4.72), TSST task (31.00 ± 5.57), and after N-back task (31.48 ± 8.18), *p*’s > 0.05. In contrast, for the stress group, participants’ anxiety levels were elevated after the TSST task (45.71 ± 11.53), compared to the baseline(32.68 ± 5.96), *p* < .001. The anxiety score after the TSST task was also significantly higher than that after the N-back task (35.33 ± 9.65), *p* < 0.001. No significant difference was found between baseline and after N-back task anxiety levels, *p* = 0.931.

### Heart rate

The change in heart rate relative to baseline is shown in Fig 3 (b). There was a significant main effect of group (*F*(1, 40) = 9.08 *p* = 0.004, *η* _*p*_^2^ = 0.19). Heart rate was significantly higher in the stress group than in the control group. The main effect of time was also significant (*F* (2, 80) *=*78.20, *p* < 0.001, *η* _*p*_ ^2^ = 0.66). The heart rate of participants was significantly higher after the TSST than it was at baseline or after the N-back tasks. The interaction between group and time was also significant (*F*(2, 80) = 39.25, *p* < 0.001, *η*_*p*_ ^2^ = 0.40).

Further analyses showed that, for the control group, no significant differences were found in heart rates between the baseline period (76.29 ± 11.23), the TSST task (79.14 ± 10.52), and after the N-back task (74.96 ± 9.56), *p*’s > 0.05. In contrast, the stress group showed a significant difference between the baseline (78.97 ± 10.65) and the post-TSST task (98.15 ± 15.66), *p* < 0.001. Additionally, there was a significant difference between the TSST task and post-N-back task (85.39 ± 13.44), *p* < 0.001. Lastly, a significant difference was found between the baseline and post-N-back task, *p* < 0.001.

### Cortisol concentrations

The change in cortisol concentrations relative to baseline is shown in Fig 3 (c). There was a significant main effect for the group (*F*(1, 40) = 150.04, *p* < 0.001, *η* _*p*_ ^2^ = 0.79). Cortisol was significantly higher in the stress group than in the control group. The main effect of time was significant (*F*(2, 80) *=*16.69 *p* < 0.001, *η* _*p*_^2^ = 0.29). Participants’ cortisol levels were significantly higher after the TSST than it was at baseline or after the N-back tasks. The interaction between group and time was also significant (*F*(2, 80) = 11.72, *p* < 0.001, *η*_*p*_^2^ = 0.23).

Further analyses showed that, for the control group, no significant differences were found in cortisol concentrations between the baseline period (8.69 ± 5.88) and the post-TSST task (9.98 ± 5.10), *p* > 0.05. but a significant difference was found between the post-TSST task and the post-N-back task(6.21 ± 2.54), *p* < 0.001. In contrast, the stress group showed a significant difference between the baseline (5.92 ± 5.06) and the post-TSST task (11.38 ± 5.18), *p* < 0.001. Additionally, there was a significant difference between the baseline and the post-N-back task (9.20 ± 6.19), *p* < 0.001.

### Behavioral data

#### Reaction time (RT)

There was a significant main effect of memory load (*F*(2, 80) = 75.49, *p* < 0.001, *η* _*p*_^2^ = 0.6). The RT in the 1-back tasks (525.67 ± 121.50 ms) was significantly longer than that in the 0-back tasks (441.64 ± 68.14 ms), and the RT in the 2-back tasks (645.86 ± 166.36 ms) was significantly longer than that in the 0-back and 1-back tasks. The main effect for the group was not significant (*F(*1, 40) = 0.69, *p* = 0.41). The interaction of the group and memory load was not significant (*F*(2, 80) = 1.43, *p* = 0.26).

#### Accuracy

The main effect of memory load was significant (*F*(2, 80) = 69.94, *p* < 0.001, *η* _*p*_ ^2^ = 0.64). Accuracy in the 0-back tasks (0.96 ± 0.03) was significantly higher than in the 1-back (0.93 ± 0.04) and 2-back tasks (0.88 ± 0.05), and it was significantly higher in 1-back tasks than in 2-back tasks. The main effect for the group was not significant (*F*(1, 40) = 1.04, *p* = 0.31). The interaction of the group and memory load was significant (*F*(2, 80) = 3.15, *p* = 0.048, *η* _*p*_^2^ = 0.07). Further analysis revealed that the stress group had a significantly higher accuracy (0.95 ± 0.02) than the control group (0.92 ± 0.02) in the 1-back task (*p* = 0.03, *η*_*p*_ ^2^ = 0.11).

## Discussion

The present study focused on the effects of acute stress on pilots’ WM. Stress induction via a modified Trier Social Stress Test (TSST) was successful, as evidenced by increased self-reported anxiety, an accelerated heart rate, and elevated cortisol levels in the stress group. As expected, increasing memory load was associated with slower reaction times and decreasing accuracy, consistent with findings from previous studies.

### Working memory performance under acute stress

The results of the WM test showed that as the memory load increased, the participants’ reaction time increased, and accuracy showed a gradual decrease. This is consistent with the results of most previous studies, where the strain on WM was elevated when the memory load in the N-back task was increased [17]. Studies that have used EEG techniques found that the EEG components associated with WM updating were impaired at high-load levels in participants under acute stress [18], which may help explain the decreased performance on the N-back task. No significant differences were observed in reaction times between the stress and control groups across memory load conditions, but accuracy was notably higher in the stress group under a moderate load. These findings support the emotion regulation hypothesis, suggesting that acute stress enhances pilots’ accuracy in a task with a moderate memory load. The implications of these results are crucial for professions such as aviation, where managing stress and maintaining optimal WM performance are key to safety. An improved understanding of the effect of acute stress on WM can inform stress management training development, potentially bolstering pilots’ performance under stress.

### The modulatory role of memory load in acute stress effect on working memory

Memory load also moderates the impact of acute stress on working memory, restricting the generalizability of our findings. For instance, Cornelisse (2011) et al. found that acute stress could increase WM levels in a group of male participants under moderate memory load conditions [12]. Specifically, male participants had faster reaction times in the moderate memory load condition after completing the TSST. Although there was no significant increase in accuracy, the performance of the stress group was better than that of the control group. Duncko et al. (2009) similarly found that the stress group had significantly faster reaction times in the high cognitive load condition than the control group [13]. When reviewing the controversial results regarding the effects of acute stress on WM under different memory loads, one researcher found that acute stress impaired WM under high memory load conditions [14]. Some researchers also found that stress impaired cognitive tasks only when the memory load was low [15]. We speculate that three main causes are responsible for this.

First, methods of inducing acute stress can vary (psychological, physiological, and pharmacological), leading to different intensities and durations of stress [19, 20], which may affect working memory performance. In this study, we used a psychologically-based method (TSST), which may induce moderate stress compared to physical or pharmacological stressors. Pessoa et al. (2009) found that a moderate intensity of emotion enhanced cognitive task performance, while a high intensity of emotion impaired task performance [21]. Therefore, we hypothesized that moderate acute stress might produce similar results. It is worth noting that the relationship between individual stress levels and job performance follows an inverted U-shaped curve and is also influenced by the task difficulty [22]. In this study, moderate stress levels led to better job performance when performing moderately difficult tasks. Thus, the stress flight group had higher accuracy WM when completing the moderately difficult task (1-back) than the control group.

Second, participant characteristics, including sex, occupation, and mental state, can influence acute stress effects on WM [12, 18]. In this study, the participants were pilots who were extensive training, likely contributing to their superior WM abilities. Additionally, the study used a digital N-back task, where pilots are more sensitive to digital information such as radio frequency, altitude, and speed. Therefore, we assume that the pilots in this study had a different difficulty level for the memory load task than other subjects. This may explain the inconsistency with previous studies.

Third, it is worth noting that the present study focused on behavioral results from WM tasks. However, other studies have found significant differences in neurological results while observing no significant differences in behavioral outcomes [15, 23]. For example, Zhang et al. (2015) found no interaction between the group and memory load on reaction time but observed significant differences in the P3 component of EEG results between the acute stress and control groups under low memory load conditions. The inclusion of brain imaging techniques such as EEG and MRI may provide further insights into the underlying neurological mechanisms associated with acute stress and working memory.

### Limitations and future directions

There are several limitations to our study. Most importantly, the dominance of male participants in our sample restricts the generalizability of our findings, as gender differences may influence the impact of acute stress on working memory. Furthermore, while the method of stress induction successfully induced acute stress, it may not fully represent the variety and intensity of stress sources that pilots face in real-world tasks. In other words, the modified TSST may not mirror some typical stress in the aviation context, such as regulation and safety requirements. Additionally, our assessment of working memory performance, focused on the N-back task, offers only a singular perspective on a complex cognitive function. The measurement could be further investigated through other tasks in the future.

Future research should include more diverse samples to enhance validity. Additionally, studies could employ alternative methods for stress induction that more closely simulate the stressors encountered by pilots in their profession.

## Conclusion

In conclusion, our study presents new insights into the relationship between acute stress and working memory performance in pilots, emphasizing the moderating role of memory load. In accordance with the Yerkes-Dodson law and the emotion regulation hypothesis, we discovered that moderate stress levels induced by the modified TSST enhance accuracy under a moderate memory load. This enhancement of memory function under moderate stress provides critical insights for the aviation industry, particularly considering the pressing requirement for pilots to precisely recall and apply SOPs during unexpected circumstances, thereby promoting flight safety.

## Acknowledgments

This study was supported by the Key Laboratory of Flight Techniques and Flight Safety, CAAC (FZ2021ZZ02), the Youth Project of Humanities and Social Sciences Financed by the Ministry of Education of China (21YJC190012, 22YJC190020), and the Fundamental Research Funds for the Central Universities (J2022-005, ZJ2020-02), and the National Natural Science Foundation of China (U2133209).

